# Quantifying the regulatory effect size of cis-acting genetic variation using allelic fold change

**DOI:** 10.1101/078717

**Authors:** Pejman Mohammadi, Stephane E Castel, Andrew A Brown, Tuuli Lappalainen

## Abstract

Mapping *cis*-acting expression quantitative trait loci (*cis*-eQTL) has become a popular approach for characterizing proximal genetic regulatory variants. However, measures used for quantifying the effect size of *cis*-eQTLs have been inconsistent and poorly defined. In this paper, we describe log allelic fold change (aFC) as a biologically interpretable and mathematically convenient unit that represents the magnitude of expression change associated with a given genetic variant. This measure is mathematically independent from expression level and allele frequency, applicable to multi-allelic variants, and generalizable to multiple independent variants. We provide tools and guidelines for estimating aFC from eQTL and allelic expression data sets, and apply it to GTEx data. We show that aFC estimates independently derived from eQTL and allelic expression data are highly consistent, and identify technical and biological correlates of eQTL effect size. We generalize aFC to analyze genes with two eQTLs in GTEx, and show that in nearly all cases these eQTLs are independent in their regulatory activity. In summary, aFC is a solid measure of *cis*-regulatory effect size that allows quantitative interpretation of cellular regulatory events from population data, and it is a valuable approach for investigating novel aspects of eQTL data sets.

## Introduction

Non-coding genetic variation affecting gene regulation and other cellular phenotypes has a key role in phenotypic variation and disease susceptibility (Albert and Kruglyak 2015). One of the most commonly used methods to characterize genetic variants that affect gene expression is eQTL mapping (Schadt et al. 2003; Lappalainen et al. 2013; GTEx Consortium 2015), which identifies genetic loci where genotypes of genetic variants are significantly associated to gene expression in a population sample. Genes and variants with significant associations are often called eGenes and eVariants, respectively, and the eVariant with the best p-value in a given locus usually used as the proxy for the causal variant. The association between genotype and gene expression is typically tested by regressing gene expression on the number of alternative alleles using a linear model, and the significance of the regression slope is used to measure significance of the eQTL (Shabalin 2012; Ongen et al. 2016). eQTLs can occur either in *trans* through altering diffusible factors that affect gene expression distally or in *cis* through allelic, physical interactions on the same chromosome typically less than 1 Mb away from the eGene, which are the focus of this study. The allelic effect of *cis*-regulation leads to unequal expression of the two haplotypes (allelic imbalance) in individuals that are heterozygous for a *cis*-acting eVariant (Fig. 1A).

**Figure 1.**
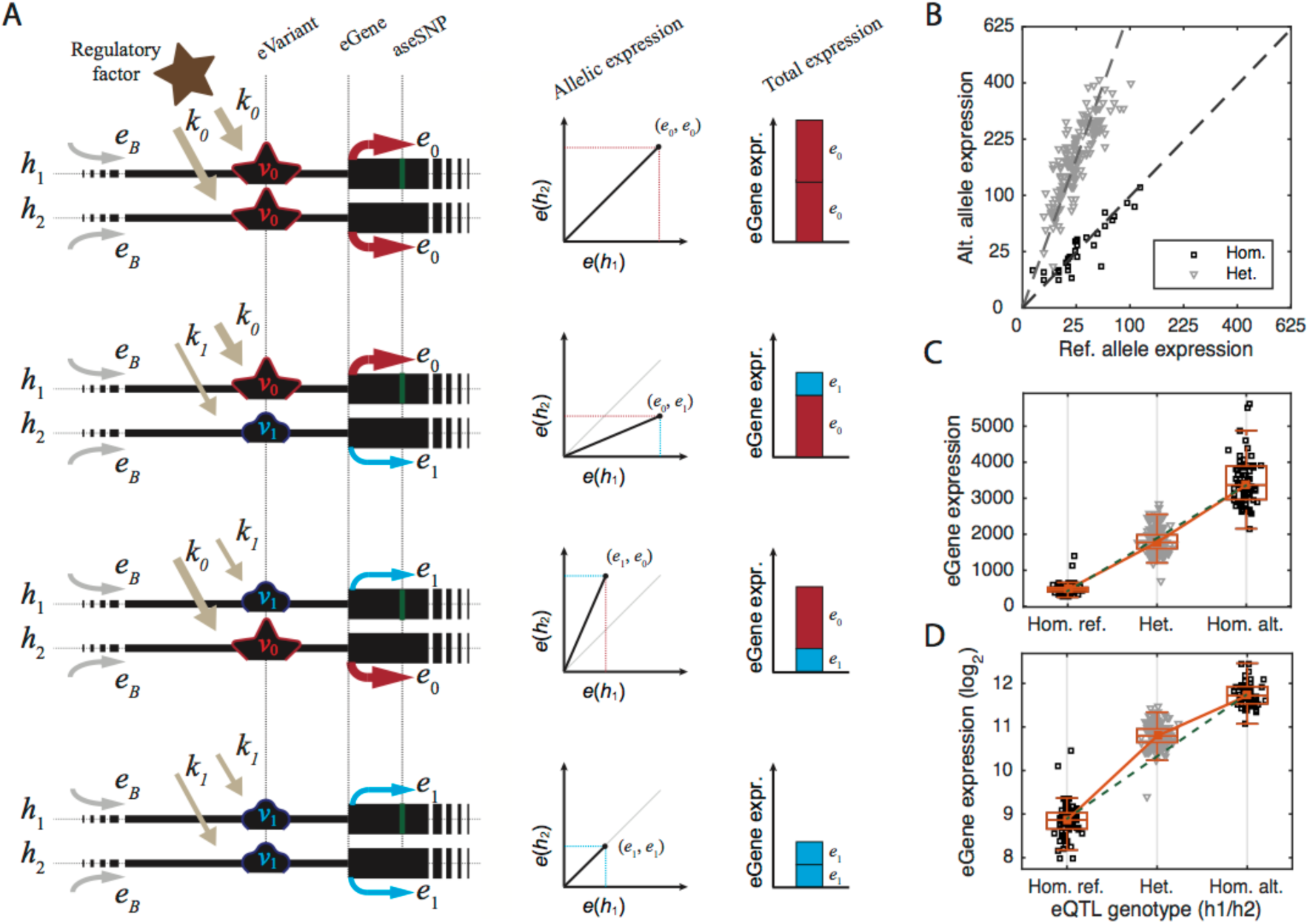
A) Schematic representation of *cis*-regulatory eQTL model in Eq. 1, 2. B) Example of allelic expression (eVariant chr5:96252589 T/C; eGene *ERAP2*) in GTEx Adipose Subcutaneous. C-D) eGene expression for the same example eQTL. The green dashed line connects the median expression of the two homozygous classes. Expression is linear with number of alternative alleles (C), but the linearity is lost after log-transformation (D).

The effect size of an eQTL describes the magnitude of the effect that it has on gene expression and is an important statistic for characterizing the nature of regulatory variants. Estimating the relative effect of eQTL alleles on expression levels has applications in computational functional genetics analysis, as well as in analysis of genetic regulatory variants by experimental assays such as genome editing (Tewhey et al. 2016; Ulirsch et al. 2016; Vockley et al. 2015; Arnold et al. 2013; Canver et al. 2015; Wright and Sanjana 2016). However, thus far there has been no consensus definition for eQTL effect size, let alone one that allows a direct biological interpretation, with previous eQTL studies using different units and approaches. The most widely used measure of effect size is simply the regression slope, a readily available statistic from eQTL calling tools (Shabalin 2012; Gutierrez-Arcelus et al. 2013; Tung et al. 2015; Lee et al. 2015). Other statistics include slope of linear regression based on log-transformed expression (Flutre et al. 2013; Battle et al. 2014), and estimation of the difference between genotype classes, such as the mean difference in expression between heterozygous and the more common homozygote class, sometimes with log transformation or scaling by mean (Gutierrez-Arcelus et al. 2015; Josephs et al. 2015). The proportion of expression variance in the population explained by an eQTL is a widely used statistic that is informative of population variance but not of the molecular effect of an eQTL (Wright et al. 2014; Kirsten et al. 2015; Grundberg et al. 2012). A recent method, developed simultaneously and independently from our work, uses the ratio between the slope and intercept of the linear regression in a variance stabilized model (Palowitch et al. 2016). While all these approaches provide estimates that are generally correlated with *cis*-regulatory effect of a given variant, they often lack a well-defined unit that enables biological interpretation of the effect size. Furthermore, many of these statistics are confounded by nuisance variables such as genotype frequency, gene expression level or technical or environmental variation. These limitations can confound downstream analysis.

*cis*-acting regulatory variation is known to be reflected in both allele-specific expression (ASE), and total gene expression data as incorporated in previous statically involved *cis*-eQTL calling methods (Pickrell et al. 2010; Sun 2012; van de Geijn et al. 2015; Hu et al. 2015; Kumasaka et al. 2016). In this study, based upon the mechanistically justified model of additive *cis* genetic effects on gene expression, we define the log-ratio between the expression of the haplotype carrying the alternative allele to the one carrying the reference allele, the log *allelic fold change* (aFC), as a biologically interpretable and mathematically convenient measure of *cis-*regulatory effect size. We provide a thorough description of the derivation and properties of this measure, including its generalizations that enable analysis of multi-allelic genetic variants and joint modeling of multiple *cis*-regulatory variants. We make the calculation of eQTL effect sizes accessible to the wide eQTL community by practical guidelines and tools, and provide effect sizes for all *cis*-eQTLs in the GTEx data set (co-submitted -(Aguet et al. 2016)). We characterize the empirical trends across the effect sizes in GTEx data, demonstrating a good fit between the empirical results and simulations, and describing factors correlated with observed allelic fold changes. Finally, we demonstrate application of *cis*-regulatory model extended for joint analysis of allelic fold changes in eGenes with two eQTLs in GTEx data.

## Results

### 1. Model

#### 1.1 Additive model of regulation

For a given gene and a given *cis*-regulatory variant, *v*, with two alleles in population, *v*_0_ and *v*_1_, we define allelic expressions *e*_0_, and *e*_1_ as the amount of transcript produced from the gene when it is located on the same chromosome copy as alleles *v*_0_, and *v*_1_, respectively. We assume that the allelic expression is determined by a shared basal expression of the gene, *e*_*B*_, driven by the cellular regulatory environment, and allele-specific factors *k*_0_, *k*_1_ ≥0, that represent distinctive effect of the allele *v*_0_, and *v*_1_ on transcription, respectively (Fig. 1A):

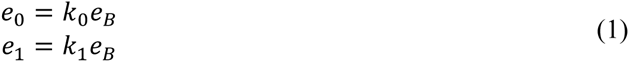

Under the *cis*-regulatory model, the regulatory effect of an allele does not depend on the genotype on the other chromosome copy, and *e*_*i,j*_, the total expression of the gene in an individual with alleles *v*_*i*_ and *v*_*j*_ on the first and second haplotype is

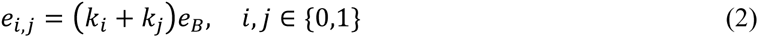

However, observational population data generally includes only relative expression quantifications. Using δ_i,j_=*k*_i_/*k*_j_ in Eq.1, the expression of haplotype carrying the alternative allele *v*_1_ is given as

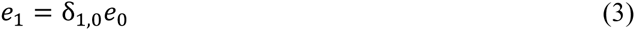

relative to *e*_0_, the expression of the haplotype carrying the reference allele. Similarly, the total relative expression of the gene is

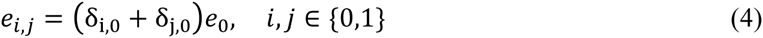

For a given *cis*-acting eVariant, we define log allelic fold change, *s*_1,0_ = log_2_ δ_1,0_, as the relative *cis*-regulatory strength of the allele *v*_1_ versus the reference allele *v*_0_. This quantity is similar to the widely used log expression fold change of differentially expressed genes, but defined between two alleles of a genetic variant. Allelic fold change of a biallelic eVariant can be directly quantified from allelic gene expression in heterozygous individuals (Fig. 1A-B; Box 1), or from summary statistics of standard eQTL linear regression between genotypes and total expression levels (Fig. 1C; Box 2). A further challenge in eQTL effect size estimation is the heteroscedasticity of noise in expression data, which violates the data normality assumptions of linear regression. Although different RNA measurement platforms such as RNA-sequencing, microarrays and other techniques have distinct technical variation profiles, biological variation in gene expression data is generally considered to be log-normally distributed (Tu et al. 2002; Whitehead and Crawford 2006; Anders and Huber 2010). However, after the commonly used variance stabilization by log transformation, gene expression is no longer a linear function, and as such cannot be solved efficiently (Fig. 1D; **Methods**). Thus, we introduce an efficient heuristic method to estimate allelic fold change from log-transformed gene expression data in linear time (Box 3). The method generates a set of four candidate aFC estimates: The first three estimates are calculated by using only two out of the three eQTL genotype classes at a time. The fourth estimate is derived using loglinear regression, utilizing the fact that log-transformed eQTL data approaches a linear function in weak eQTLs as log allelic fold change goes to zero (*s*_1,0_ → 0; **Methods**). The candidate aFC that minimizes the residual variance in log-transformed data is reported as the final estimate (**Methods**).

##### Box 1: Calculating aFC from allelic expression data.

###### Input

- Allelic expression in *N* individuals heterozygous for the top eVariant of an eQTL of interest: (*c*_0,1_, *c*_1,1_)… (*c*_0,N_, *c*_1,N_)

1. Get median ratio of the allelic counts:

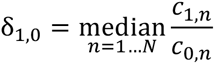

where (*c*_0,n_, *c*_1,n_) are allelic counts from the 1^st^, and 2^nd^ haplotype in the *n*^th^ individual.

###### Output

Report effect size: *s*_1,0_=log_2_ δ_1,0_

##### Box 2: Calculating aFC from gene expression data (see **Methods** for derivations).

###### Input

- eGene expression in *N* individuals: *y*_1_… *y*_N,_, where y_n_ ∈ [0, +∞)
- Number of alternative alleles in each individual: *t*_1_… *t*_N_, where t_n_ ∈ {0,1,2}

1. Use simple linear regression to model expression as a function of *t*_n_:

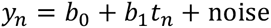
2. Use the slope *b*_1_ and intercept *b*_0_ to calculate:

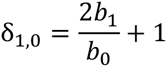

###### Output

Report effect size: *s*_1,0_=log_2_ δ_1,0_

##### Box 3: Linear time algorithm for estimating aFC from log-transformed gene expression data (see **Methods** for derivations).

###### Input

- eGene expression in *N* individuals in log_2_ scale: *z*_1_… *z*_N,_, where *z*_n_ ∈ [−∞, +∞)
- Number of alternative alleles in each individual: *t*_1_… *t*_N_, where *t*_n_ ∈ {0,1,2}

1. Calculate *m*_0_, *m*_1_, *m*_2_ as geometric mean of expression for individuals with *t*_*n*_= 0, 1, and 2, respectively.
2. Calculate the following three candidate estimates:

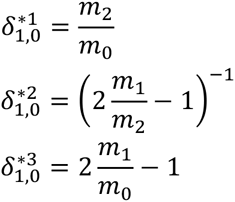
3. Use simple linear regression to model log_2_ expression as a function of *t*_*n*_:

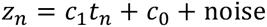
4. Use the slope *c*_*1*_ times two as the fourth candidate estimate:

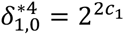
5. Use each of the four estimates 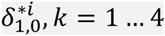 to calculate:

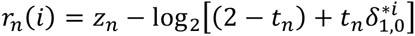 Where 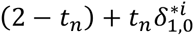 is predicted gene expression in *n*^th^ individual using the *i*^th^ estimate.
6. Pick the estimate that provides the lowest variance in the residuals:

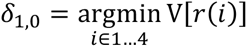

###### Output

Report effect size: *s*_1,0_=log_2_ δ_1,0_

#### 1.2 Generalization to multiple eVariants with multiple alleles

Beside clear biological interpretation, log allelic fold change has several convenient mathematical properties that facilitate downstream analysis of the values (Box 4, **Supplemental methods**), and allow generalization to analysis of multi-allelic genetic variants, as well as to joint analysis of multiple independent eQTLs for the same eGene. Here we consider the case of *N* eVariants, *v*1,…, *v*n,… *v*N acting on the same eGene independently with *m*_1_,… *m*_n_,… *m*_N_ alleles, respectively. Let 〈*i*_1_… *i*_n_… *i*_N_〉 denote a haplotype carrying the *i*_n_-th allele of the *v*n, the relative expression on this haplotype is:

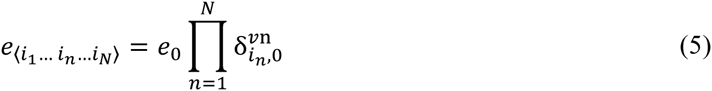

##### Box 4: Mathematical properties of log aFC as a relative measure of *cis*-regulatory effect size (see **Supplemental methods** for proofs).

1. Zero log aFC indicates the absence of regulatory difference: *s*_*i,i*_ = 0
2. Choice of reference allele only affects the sign of log aFC: *s*_*i,j*_ = −*s*_*j,i*_
3. Log aFC is additive:

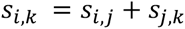
4. Log aFC associated with joint effect of independent regulatory variants, *v*1…*v*N is sum of their individual aFCs:

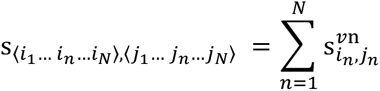 Where 〈*i*_1_… *i*_*n*_… *i*_*N*_〉 and 〈*j*_1_… *j*_*n*_… *j*_*N*_〉 are the set of present alleles on each of the haplotypes.
5. Absolute value of log aFC, *d*_*i,j*_ = |*s*_*i,j*_|, is a pseudo-metric:

i. *d*_*i,j*_ ≥ 0
ii. *d*_*i,i*_ = 0
iii. *d*_*i,j*_ = *d*_*j,i*_
iv. *d*_*i,k*_ ≤ *d*_*i,j*_ + *d*_*j,k*_

Where 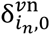 denotes the allelic fold change associated with allele *i*_n_, at the *n*^th^ eVariant *v*n versus its reference allele 0, *e*_0_ is the reference expression associated with the case, *e*_〈0… 0… 0〉_, where the haplotype carries reference alleles for all eVariants. Thus the log allelic fold difference between two haplotypes 〈*i*_1_… *i*_*n*_… *i*_*N*_〉 and 〈*j*_1_… *j*_*n*_… *j*_*N*_〉 is

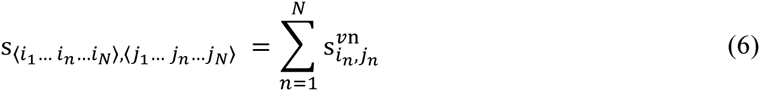

Where 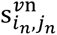 denotes the log allelic fold change associated with two, alleles *i*_n_ and *j*_n_, at the *n*^th^ eVariant. The total expression of the eGene given the genotype is

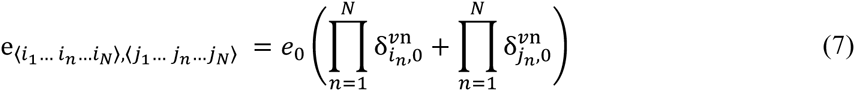

Following the *cis*-regulatory model, this inherently takes into account the independent expression of the two haplotypes according to the alleles that they carry, which is different from a simple additive model of multiple eQTLs that ignores their haplotype configuration. The last two equations can be used to simultaneously estimate effect sizes of *N* eVariants from allelic expression or transcription profiles of genotyped individuals, respectively.

#### 2. Simulation without noise

Regression slope is probably the most common measure used for estimating *cis*-eQTL effect size. However, compared to aFC, it lacks a clear biological interpretation, and is prone to systematical biases introduced by expression level and allele frequency. We demonstrate this by simulation of *cis*-eQTLs without noise (Eq. 3, 4), and comparing estimates of effect size by log aFC (Box 2, 3; since there is no noise, both methods yield identical results) linear regression slope (***b***_1_ in Box 2), and regression slope after log-transformation of the expression data (***c***_1_ in Box 3). We consider two eQTLs: one with four times higher expression of the alternative than the reference allele and another the opposite. The three measures of effect size were calculated for a fixed reference allele frequency of 50% and varying gene expression levels (Fig. 2A), and for a fixed gene expression level and varying allele frequency (Fig. 2B). Our results show that linear expression slope varies with gene expression levels, and the loglinear slope varies with allele frequency, and neither provides a quantitative estimate of the four-fold expression difference between alleles. The log aFC estimate remains insensitive to both confounding factors, and yields the correct estimate of the eQTL effect size.

**Figure 2:**
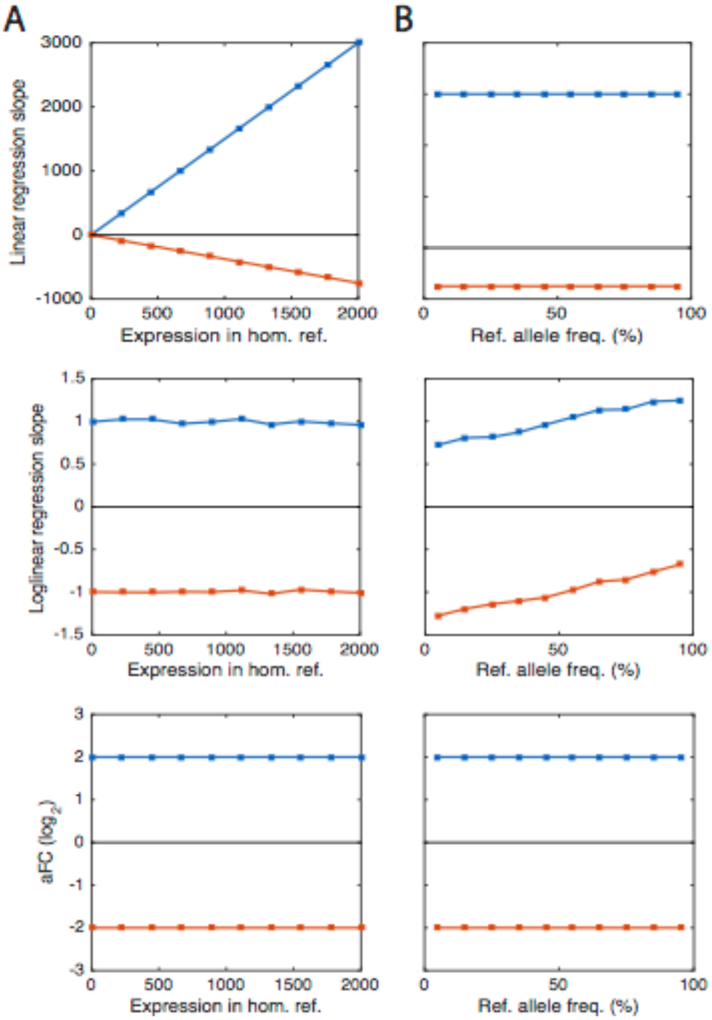
Comparison of three eQTL effect size measurements in simulations without noise. Log aFC is compared to linear and loglinear regression slopes for simulated eVariants with the alternative allele expressed four times higher compared to the reference (blue) and vice versa (red), for varying expression levels of the homozygous reference genotype (A) and varying allele frequencies (B). The results demonstrate that log aFC quantifies the biological expression difference between alleles, and is robust to changing allele frequencies and expression levels.

#### 3. Noise distribution in eQTL data and simulation with realistic noise

Next, we used simulation to evaluate how our three alternative methods for calculating aFC perform under a realistic expression noise level: M1) Linear method that uses linear regression coefficients from eQTL data as benchmark for speed (Box 2); M2) Nonlinear method that directly solves the regression problem in Eq. 17 using a standard nonlinear least square optimization tool (**Methods**) as a benchmark for accuracy; M3) Nonlinear approximation that solves the nonlinear regression problem from Eq. 17 using our heuristic solution (Box 3, Fig. 3C). In this simulation, we used simulated data of **10,000** eQTLs with varying allele frequencies and effect sizes (Eq. 3, 4), with noise added to the expression levels at **40%** coefficient of variation within genotype groups (log_10_ *ε*_*n*_ ∼ *norm*[0, *σ* = 0.17], Eq. 17) similar to what is observed in real data from GTEx (Supplementary Fig. 1). We found that at this level of noise all three methods provide highly accurate and similar estimates (Fig. 3). All estimates, especially the linear method (M1), deteriorate in eQTLs in which lower expressed allele has also a low frequency (Fig. 3B). This problem is inherent to *cis*-eQTL data and is expected to occur regardless of the expression measurement platform. Overall, the aFC estimates from the nonlinear model (M2) provided the lowest root mean squared deviation (RMSD) from the true values, with the its approximation (M3) providing only 10% worse RMSD than the nonlinear model at 1.8 times the runtime of the linear model. The linear model was 84 times faster than the nonlinear model but provided 64% higher RMSD.

**Figure 3:**
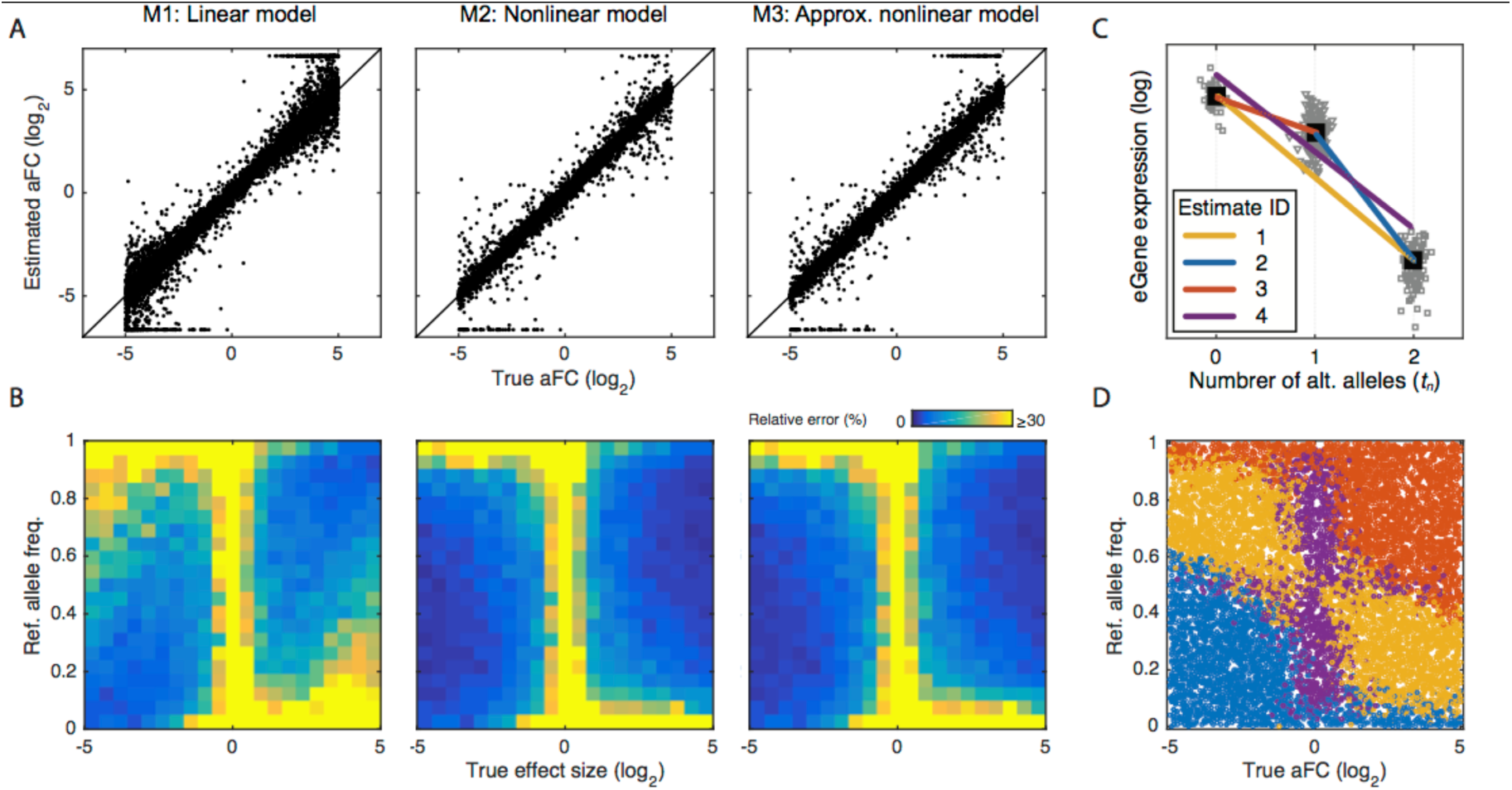
Comparison of the aFC estimation methods using simulated data. We simulated 10,000 eQTLs with noise (40% coefficient of variation), and uniformly selected log_2_ aFC (range: [-5,5]), and reference allele frequency (range: [0,1]). A) True aFC used in simulation versus identified values using linear model (M1), nonlinear model (M2), and the nonlinear model approximation (M3). At this level of noise M2 performed the best, with M1 and M3 having RMSDs of 164% and 110% of M2. B) Quality of the effect size estimates as a function of allele frequency and the true effect size, evaluated by average error relative to the true log_2_ aFC. All three estimates and particularly M1 deteriorate when the lower expressed allele is the minor allele. C-D) Schematic representation of the nonlinear model approximation method (Box 3), based on four different candidate estimates (C), and the selected estimate with minimum residual variance for each simulated eQTL as a function of reference allele frequency and the true aFC (D).

#### 4. Application to GTEx eQTLs

Next, we applied our methods for effect size estimation to the *cis*-eQTLs discovered in the Genotype Tissue Expression (GTEx) (GTEx Consortium 2013; 2015) v6p dataset, with eQTL data from 44 tissues (70-361 individuals per tissue; co-submitted (Aguet et al. 2016)), calculating aFC for all the reported eQTLs in each tissue, using the eVariant with the best p-value for each eGene. Allelic fold changes were estimated from both allele-specific expression (ASE; Box 1) and eQTL data (Box 2–3). For ASE data, we used haplotypic expression at eGenes calculated by summing allelic expression from all phased heterozygous SNPs within the gene. aFC was reported for an average of 57% of eGenes per tissue, requiring haplotypic coverage of at least 10 reads in at least 5 individuals (co-submitted (Aguet et al. 2016)). For eQTL-based aFC estimates, we log transformed normalized read counts, and corrected for confounding factors identified using PEER (Stegle et al. 2012) and the top three principal components of the genotype matrix (see methods, Eqs. 23–24). The log aFCs for the eQTLs were calculated using the three models as in the simulation study, and constrained to ±log 100. All three eQTL methods provided highly similar aFC estimates with high concordance to ASE-based estimates (Fig. 4A, C). The effect sizes were more discordant between ASE and eQTL-based estimates when the rare allele was the lower expressed allele, as predicted by the simulation study (Fig. 4B). The nonlinear model provided the best estimates as evaluated by RMSD from ASE-based estimates, and was closely trailed by the nonlinear approximation method (Fig. 4C). Thus, for the rest of the analyses we used only the nonlinear approximate method as it provided both high accuracy and speed. Finally, we tested the effect of quantile normalization that enforces log-normality of expression data within each genotype. While this is commonly used to avoid outlier effects, we did not observe improvement of the effect size estimates (Fig. 4D).

**Figure 4:**
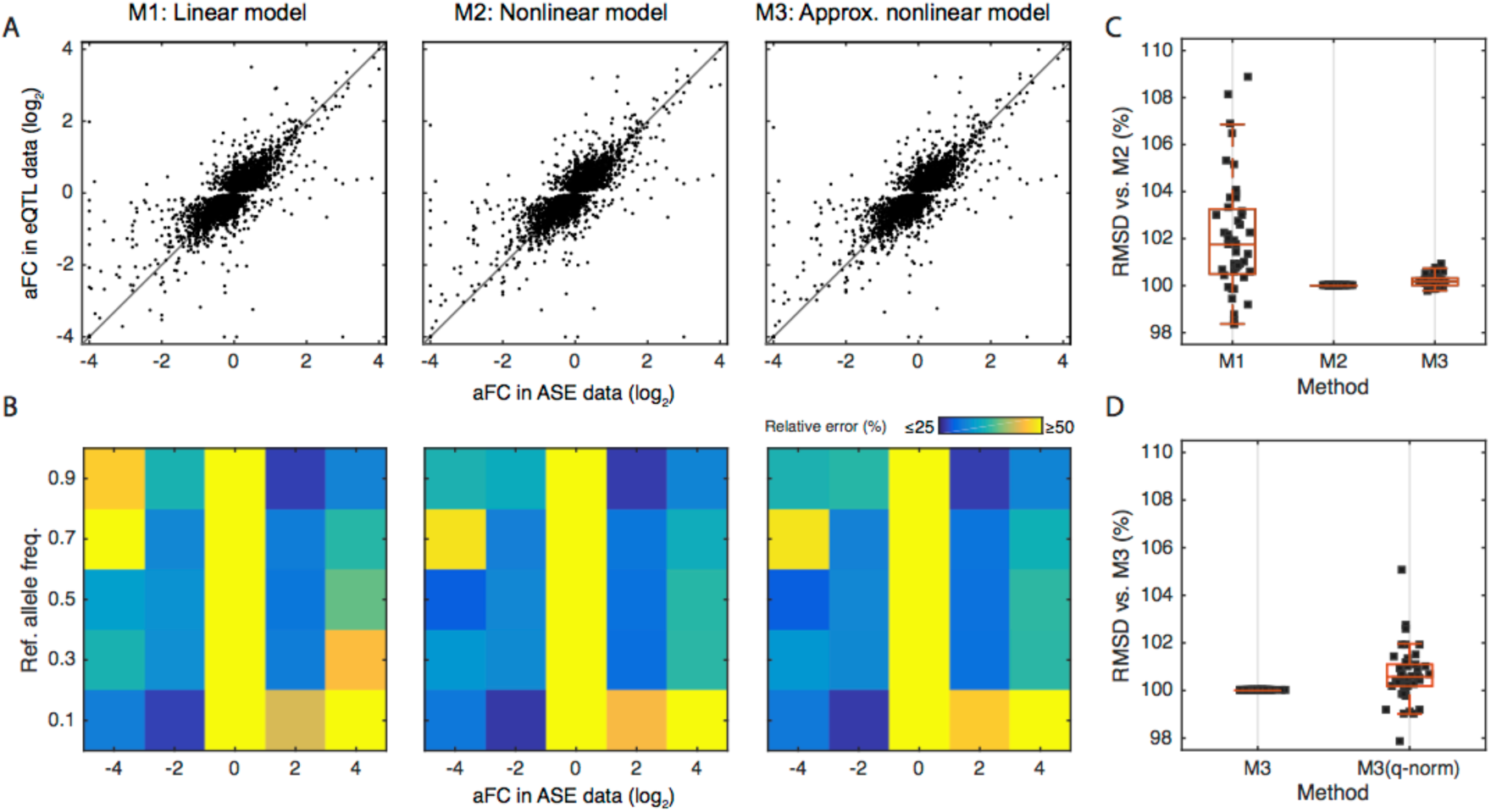
Comparison of the methods for estimating aFC using GTEx data. A) Allelic fold change as estimated from ASE data versus estimates from eQTL data using linear model (M1), nonlinear model (M2), and the nonlinear model approximation (M3) for all top eQTLs in Adipose Subcutaneous. All three estimates are ∼75% correlated with estimates form ASE data. B) Quality of the eQTL estimates as a function of allele frequency and the aFC estimate from allelic expression data, evaluated by average relative error between aFC from ASE data and from eQTL estimates. C) Concordance between the estimates from allelic expression and eQTL data as evaluated by RMSD between the most accurate method, M2, and the other two methods. Each dot represents one tissue in GTEx. D) Concordance between the estimates from ASE and eQTL data as evaluated by RMSD, comparing M3 to M3 applied after quantile normalization within each genotype group. Each dot represents one tissue in GTEx.

The empirical distributions of aFCs for eQTLs detected in different GTEx tissues are highly dependent on the sample size, since tissues with lower sample size lack power to detect weak eQTLs (Fig. 5A). The effect size estimates from eQTL and ASE data are highly similar, but on average 1.45% (CI: [1.3, 1.6]) smaller across the tissues when estimated from ASE data (Fig. 5B, C). This mild overestimation of the effect size involving weaker eQTLs is consistent with potential winner’s curse in the eQTL calling stage (Fig. 5D). This highlights the added value of ASE-based estimates alongside eQTL data. We next analyzed the correlation of aFC with other properties of the eVariant or eGene. Low-frequency eVariants tend to have higher effect sizes (Fig. 5E), likely due to differences in power to detect eQTLs and other statistical artifacts. eGenes with high expression levels, expression in multiple tissues, and high coding region conservation measured by RVIS (Petrovski et al. 2013) have lower effect sizes (Fig. 5F-H), which suggests that genes under strong selective constraint are less likely to tolerate regulatory variants with high effect sizes. Further biological interpretation of effect sizes across eVariants in different annotations and eGenes of different biotypes and eQTLs that are tissue-specific or shared is described in ((Aguet et al. 2016); co-submitted). In these and other downstream analyses of eQTL effect sizes, it is important to correct for correlated factors such as sample size and allele frequency. Even though our simulations demonstrate that aFC is highly robust to key confounders, differences in the power of eQTL mapping will always affect the properties of discovered eQTLs, including effect size distribution.

**Figure 5:**
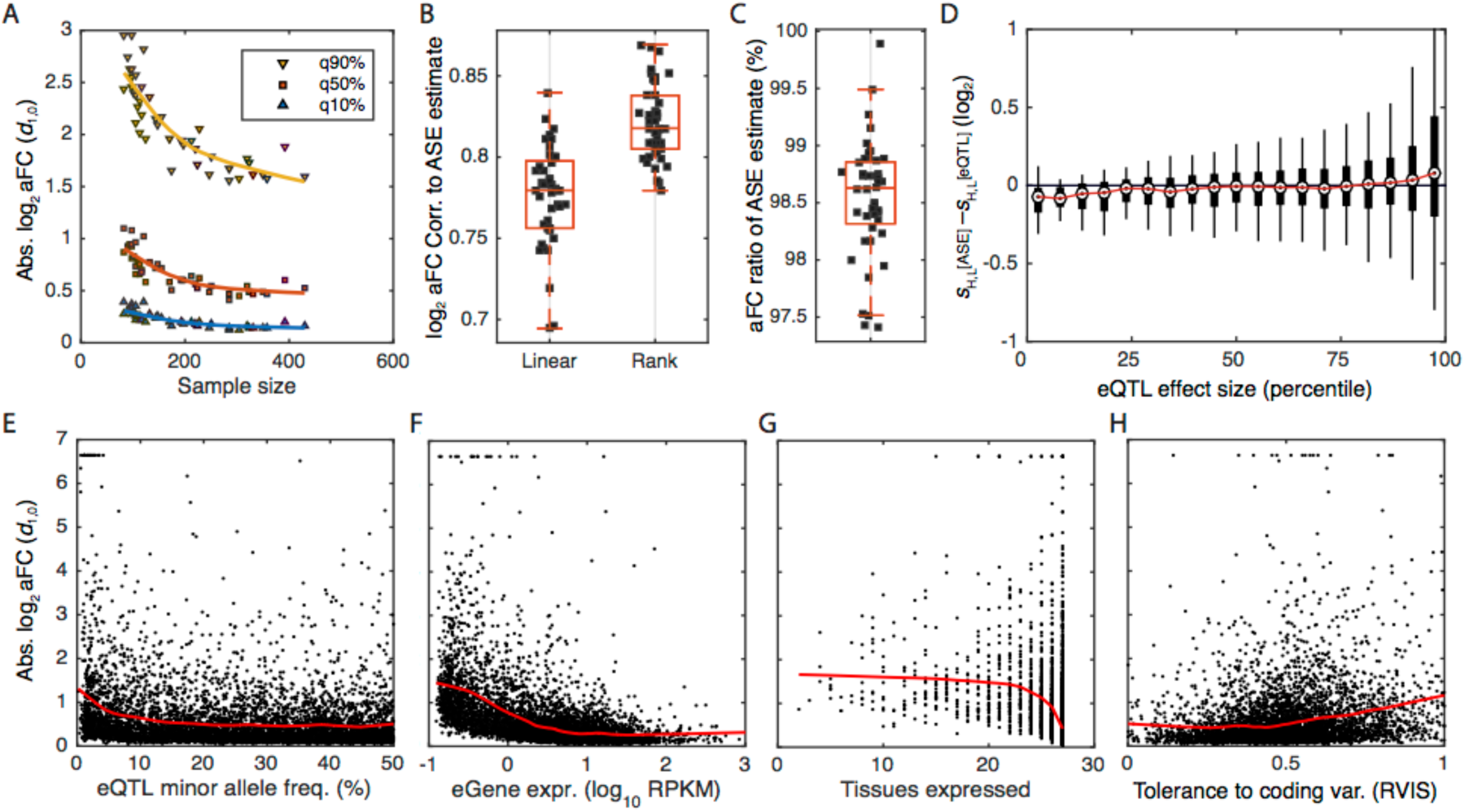
Empirical properties of the aFC distributions in GTEx data. All aFC values are calculated with the nonlinear approximation method (M3). A) Distribution of absolute log_2_ aFC across tissues as a function of sample size. Each point represents a tissue in GTEx data, and 90%, 50%, and 10% quantiles of absolute aFC across a tissue are shown. B-C) Correlation of log_2_ aFC estimates (B), and the median ratio of aFC estimates (C), derived from eQTL and ASE data. Each point corresponds to one GTEx tissue. D) Difference between the aFC estimates from allelic expression (*s*^ASE^), and eQTL (*s*^eQTL^), as a function of absolute average aFC (|*s*^ASE^ + *s*^eQTL^|/2), with H, and L referring to higher and lower expressed alleles of each eQTL in Adipose Subcutaneous, respectively. Estimated effect size form ASE data tend to be smaller in weak eQTLs and larger for stronger eQTLs as compared to those derived using eQTL data. E-H) Distribution of absolute log_2_ aFCs calculated from GTEx Adipose Subcutaneous as function of minor allele frequency (E), gene expression level (F), number of tissues where the gene is expressed >0.1 RPKM in ≥10 individuals (G), and logistic-transformed RVIS, a measure of each gene’s tolerance to variation in the coding region ((Petrovski et al. 2013); H). Red line shows fit by robust locally weighted scatterplot smoothing.

The allelic fold changes of GTEx eQTLs are provided in the GTEx portal (see Data Access section). Additionally, we implemented the linear model (M1) and the nonlinear approximation model (M3) in a python script (see **Data Access**) that takes as input the standard file formats used also by the FastQTL software for eQTL calling. This makes calculation of aFC for other eQTL datasets straightforward and fast.

#### 5. Independent eQTLs in GTEx

Iterative greedy procedures have been utilized to find multiple independent eQTLs signals for each eGene in the GTEx data (co-submitted (Aguet et al. 2016)). We used GTEx eGenes with two independent eQTLs to demonstrate how the aFC calculation can be extended to gain mechanistic insight into more complex eQTL patterns. The expression model in Eq. 7 written for two biallelic eVariants was used in a nonlinear regression to simultaneously estimate the aFC associated with both eQTLs (Fig. 6A; **Methods**). These estimates were used to predict the relative expression of the two haplotypes between the 16 possible haplotypic combinations. We found that the predicted values from eQTL data correlate well with the observed values in ASE data across the genotypes (median *r*=0.81, Fig. 6B-D). While the used model accounts for specific arrangement of the alleles for the two eVariants on haplotypes (e_〈11〉,〈00〉_ ≠ e_〈10〉,〈10〉_, Eq. 7), it assumes that the two eVariants act independently 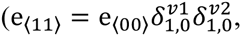 Eq. 5). In order to analyze how well the data is described assuming the independence of the two eVariants, we relaxed this assumption by defining the joint genotype of the two eVariants as the genotype of a hypothetical variant with four possible alleles. We used Eq. 7 written for one four-allelic eVariant to separately estimate the aFC associated with each of the two eVariants, and aFC of their co-occurrence. We found that the estimates from the two models generally agree very well (Fig. 6C). We used the Bayesian information criterion within a bootstrapping scheme to decide if relaxing the regulatory independence assumption provides a significantly better description of the data. This could be a sign of biological mechanisms such as epistasis or dosage compensation as well as confounding factors such as linkage disequilibrium or expression quantification artifacts (Brown et al. 2014; Hemani et al. 2014; Wood et al. 2014; Fish et al. 2016). After accounting for the increased model complexity and uncertainty associated with sampling distribution we found that only in 0.2% (range across tissues [0, 0.42]) of the two eQTLs for the same gene in GTEx data the regulatory independence model fails to provide an adequate fit (Fig. 6D, **Table S1**).

**Figure 6:**
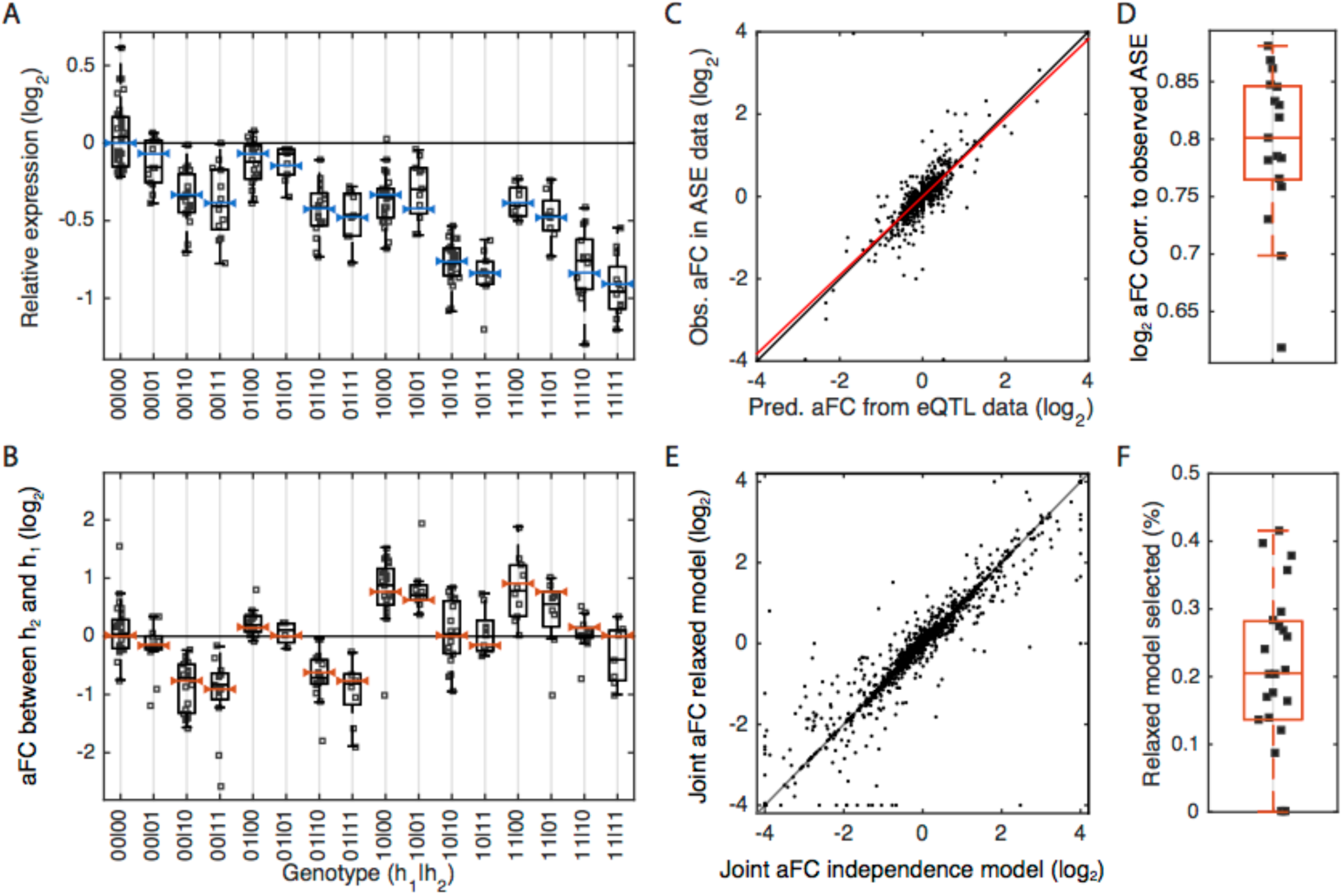
Joint analysis of aFCs for GTEx eGenes with two eQTLs. A) An example of relative expression of eGene *ZC3H3* and the model fits for different genotype groups of its two eQTLs (eVariant1: chr8:144633728 A/G and eVariant2: chr8:144556836 G/A) in GTEx Adipose Subcutaneous. The effect size of the first and the second eQTLs are −0.77 and −0.14 as measured by log_2_ aFC. Each dot represents observed expression in one individual, scaled relative to the expression at all-reference genotype. The blue bars show model fits from the two-eQTL model based on regulatory independence assumption. Reference and alternative alleles are denoted by 0 and 1, respectively, and haplotypes are separated by “|” sign (e.g. 10|11 corresponds to the cases that one haplotype carries alternative and reference alleles for eVariant1 and eVariant2, respectively, and the other haplotype carries the alternative allele of both eVariants.). B) Expression of the second haplotype relative to the first haplotype, observed in ASE data. The red bars show expected haplotype expression ratios based on the model in panel A, learned on the eQTL data. C) aFC between two haplotypes as predicted from eQTL data compared to aFC observed in ASE data for all eGenes with two eQTLs in Adipose Subcutaneous. Each dot represents one randomly selected genotype for on eGene. Red line indicates the robust linear fit (*y*=0.9*x*+0.002). D) Predicted and observed allelic fold change for all eGenes with two eQTLs, calculated from eQTL and ASE data, respectively, in each tissue with more than 200 eGenes with two eQTLs. E) *cis*-regulatory effect size associated with co-occurrence of the alternative alleles of the two eQTLs, as predicted under regulatory independence model or learned using the relaxed model. F) Percentage of the two eQTLs that are not well described using the independent regulatory assumption across all tissues with more than 200 eGenes with two eQTLs.

### Discussion

Despite over a decade of eQTL analysis and its increasingly widespread use in functional and medical genetics, eQTL effect size has lacked a clear, biologically interpretable, and computationally feasible definition. Here, we described log allelic fold change, a generalizable measure of *cis*-regulatory effect size that captures the independent regulation of haplotype expression in *cis*. Log aFC is consistent across expression levels, allele frequencies and holds mathematically convenient properties that facilitate its application for downstream analysis. aFC provides uniform estimates from both allelic expression and *cis*-eQTL data, and replication of *cis*-eQTLs using orthologous ASE data from the same samples can complement classical replication with an independent sample. While the correlation between effect sizes estimated from ASE and eQTL data is high, this is still likely an underestimate, and could be improved by using methods that produce more accurate measures of haplotypic expression (Castel et al. 2016). The two alternative aFC calculation methods provided use untransformed and log-transformed eQTL data to account for additive and multiplicative noise, respectively. We showed that the estimates that utilize log-transformed data are generally better. However, both methods perform well, and the preferred noise model can vary depending on the expression measurement platform and upstream preprocessing pipelines that have been utilized. We benchmarked aFC for RNA-sequencing data, the most popular platform for expression level quantification, but aFC is a general measure and the presented methods can be directly applied to data from other quantification platforms such as microarray and qPCR. Systematic extension of aFC-based model of *cis*-regulation to multiple alleles and multiple eQTLs, as demonstrated for the eGenes with two eQTLs in GTEx, allows investigating more complex problems while maintaining mechanistic interpretability of the results. Finally, we introduced practical guidelines and a tool for estimating aFC from real data, and provided a catalog of *cis*-eQTL effect sizes across all GTEx tissues as a resource for future studies.

A biologically interpretable and well-defined eQTL effect size estimate enables better understanding of the effects of regulatory variants at many levels. In downstream analyses of GTEx effect sizes (co-submitted (Aguet et al. 2016)), we investigate differences in effect sizes between eGene types, eVariant annotations, and eQTL tissue specificity. Even though aFC itself is unbiased with respect to allele frequency and expression level, we showed here that it is essential for all downstream analyses to take into account factors that indirectly confound the effect size distribution via eQTL discovery power. eQTL effect size quantification will be valuable for making quantitative comparisons between effects on gene expression and other phenotypes at the cellular and physiological level. Indeed, our method is generally applicable to estimating effect size of *cis*-regulatory variants affecting other cellular traits such as methylation, chromatin state, and protein levels. Furthermore, the additive nature of log aFC makes it a useful tool for characterization of variation in eQTL activity across cellular or environmental contexts in the future. For disease-associated eQTLs, understanding the relationship between the quantitative expression effect in the cells and disease risk will be important for understanding molecular mediators of disease risk. Finally, the recent development of experimental approaches such as MPRA (Tewhey et al. 2016; Ulirsch et al. 2016), STARR-seq (Vockley et al. 2015; Arnold et al. 2013) and CRISPR genome editing assays (Canver et al. 2015; Wright and Sanjana 2016) has created demand for translating summary statistics of eQTL mapping to quantifications that are interpretable as reflecting molecular events in the cell. Our biologically interpretable estimates of *cis*-eQTL effect sizes from population data can be directly compared to *in vitro* quantification of regulatory variant effects.

### Methods

#### 1. Estimating *cis*-regulatory effect of an eVariant from allelic expression data

Standard RNA sequencing reads can be used to measure the expression of each of the two gene copies, via allelic counts in individuals carrying a heterozygous SNP (aseSNP) inside the transcribed region of the gene (Castel et al. 2015). Allelic counts provide measurement of the true allelic expression *e*_0_, and *e*_1_ from Eq.1 in a given sample on a relative scale. Since both measurements are drawn from the same sample they share the same basal expression (*e*_*B*_ in Eq. 1) and thus, in absence of noise, the ratio between the two allelic counts directly reflects the effect of the *cis*-regulatory variant. Given allelic expression data from a set *N* of individuals heterozygous for an eVariant of interest, the allelic fold change can therefore be robustly estimated as

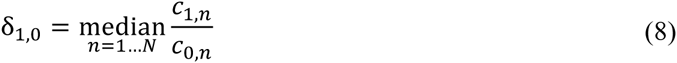

where c_0,n_ and *c*_1,n_ are the allelic counts in the *n*^th^ individual for haplotype carrying reference and alternative allele for the *cis*-regulatory variant respectively. Here we assume phasing between the regulatory alleles and the aseSNP alleles are known. In cases when phasing information is not available the magnitude of the regulatory effect size can be calculated as

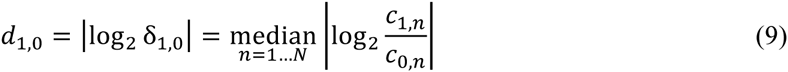

However, this estimate without phasing information is more sensitive to noise, and will systematically overestimate the effect size in cases where the true effect size is small in magnitude and the variation in allelic counts is dominated by measurement noise.

#### 2. Estimating *cis*-regulatory effect of an eVariant from gene expression data

##### 2.1 Gene expression is linear with the number of alternative alleles for biallelic eVariants

Using Eq. 4 we can derive gene expression in an individual as function of the number of alternative alleles, *t*:

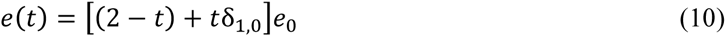

where *t* is 0,1, and 2 for individuals homozygous for reference allele, heterozygous, and homozygous for alternative allele, respectively. This equation can be written as

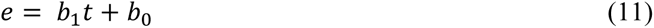

where

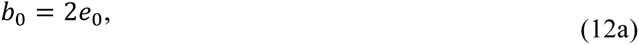

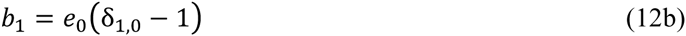

showing that total gene expression under a *cis*-regulatory model is a linear with the number of alternative alleles of the variant (Fig. 1C). For estimating the aFC from expression data we consider two cases of noise distribution, additive and multiplicative noise.

##### 2.2 Estimating aFC from eQTL data with additive noise

Under an additive noise model, the measured gene expression in the *n*^th^ individual, *y*_*n*_, is the true expression, *e*(*t*), plus a normally distributed noise, *ε*_*n*_, with zero mean and unknown variance. Using *e*(*t*) from Eq. 10:

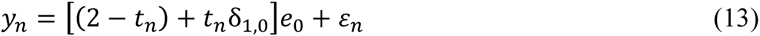

where *t*_*n*_ is the number of alternative allele in the individual. Similar to Eq. 10, Eq. 13 can be written in linear form:

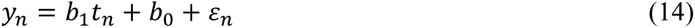

Maximum likelihood estimates for *b*_0_ and *b*_1_ can be derived efficiently using ordinary least squares, and solving Eqs. 12a and 12b, for δ_1,0_, the allelic fold change is:

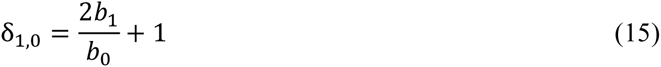

##### 2.3 Estimating aFC from eQTL data with multiplicative noise

Assuming a multiplicative noise model the measured gene expression in the *n*^th^ individual, *y*_*n*_, is the true expression, *e*(*t*), multiplied by a noise, *ε*_*n*_, such that log *ε*_*n*_ is normally distributed with zero mean and unknown variance. Substituting *e*(*t*) from Eq. 10 again:

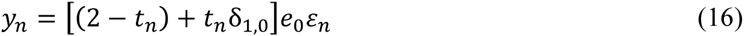

Due to the multiplicative noise, this equation can no longer be solved as a simple linear regression problem. Applying log transformation to both sides:

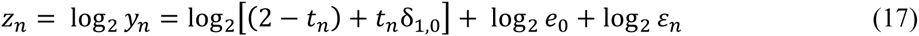

the noise is captured by log_2_ *ε*_*n*_, which is additive and normally distributed, but the right side of the equation is no longer linear with the number of alternative alleles (Fig. 1D). Using nonlinear least squares optimization Eq. 17 can be solved to derive maximum likelihood estimates for the effect size δ_1,0_ directly.

##### 2.4 Efficient approximation of aFC from eQTL data with multiplicative noise

Nonlinear least squares optimization needed for solving regression problem in Eq. 17 is done using iterative numerical optimization that is relatively slow procedure and not always straightforward to implement. In order to improve efficiency, we use four simplified linear models to derive four candidate estimates of the effect size, and choose the one that provides the highest likelihood of the data. First, we derive three estimates of the regulatory effect size using the ratio of the expressions between each of the two genotypes:

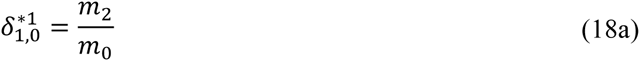

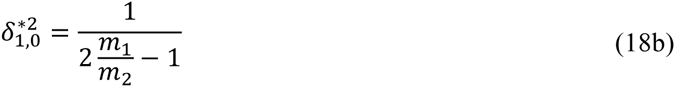

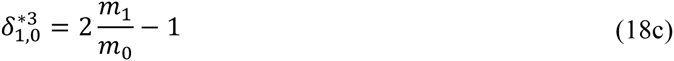

where *m*_0_, *m*_1_, and *m*_2_ are the geometric means of expression in the samples homozygous for reference allele (*t*_*n*_ = 0), heterozygous (*t*_*n*_ = 1), and homozygous for the alternative allele (*t*_*n*_ = 2) respectively (See **Supplemental methods**). When the *cis*-regulatory effect size approaches zero, the log transformed gene expression is linear with number of alternatives alleles (See **Supplemental methods**). Therefore, the nonlinear model in Eq. 17 can be well approximated with linear regression in cases where the effect size is small. We regress log-transformed expressions on the genotype:

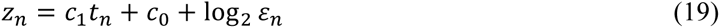

and calculate the fourth effect-size estimate as (See **Supplemental methods**)

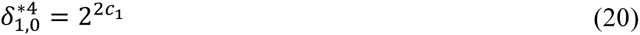

Residual of the fit, *r*_n_, in the *n*^th^ sample for a given effect size estimate, 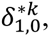 is

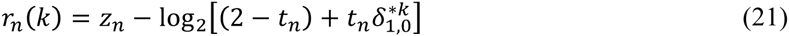

The estimate with lowest variance of the residuals among the four candidates is reported:

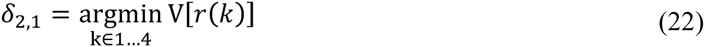

#### 3. Simulation experiment

The simulated dataset includes 200 individuals, and 10,000 eGenes each associated to exactly one eQTL. Each eQTL has two alleles, frequency of the reference allele, *f*_0_, was drawn from a uniform distribution for each eQTL (*f*_*0*_ ∼ *uniform*[0,1]). eQTL genotype in each individual was decided using two Bernoulli trials. Reference and alternative allele induce expressions *e*_0_, and *e*_1_= δ_1,0_ *e*_0_ in the eGene in *cis*-, respectively (Eqs. 1-2). The expression *e*_0_ is generated for each eGene randomly across four orders of magnitude (log_10_ *e*_0_ ∼ *uniform*[0,4]). Similarly, the aFC, δ_1,0_, was assumed to be uniformly distributed in logarithmic scale (log_2_ δ_1,0_ *∼ uniform*[-5,5]) across simulated eQTLs. In order to choose a realistic noise level we used data from all eGenes associated with eQTLs in GTEx. For each eQTL genotype class expression mean and variance of the associated eGene was calculated. As expected gene expression was highly heteroskedastic with mean-variance relationship resembling that of multiplicative noise by log-normal distribution (Fig. S1). We used average within genotype standard deviation of log_10_ transformed gene expression to add log-normal noise in the simulation (log_10_ *ε*_n_ ∼ *norm*[0, *σ* = 0.17], Eq. 17).

#### 4. Estimating aFC for GTEx eQTLs

Haplotypic counts were generated as describe in ((Aguet et al. 2016) co-submitted). Briefly, allelic counts for each sample were generated from uniquely aligned RNA-seq reads for all heterozygous SNPs from OMNI Array imputed genotypes using the GATK *ASEReadCounter* tool (Castel et al. 2015). SNPs covered by less than 8 reads, those that showed bias in mapping simulations (Panousis et al. 2014), had a UCSC 50-mer mappability lower than 1, or those without evidence for heterozygosity (Castel et al. 2015), were filtered. Haplotypic counts were generated by summing allelic counts within each gene using population phasing. For eQTL data, expression counts were scaled for the total library size, and one pseudo-count was added to smooth the normalized counts. Log transformed expression data was corrected for confounding factors identified using PEER (Stegle et al. 2012) and the three top principal components of the genotype matrix uniformly for all three tested methods: linear, nonlinear, and nonlinear approximation. The correction was done in two steps: First, log transformed expression profile of the eGene in *n*^th^ sample, *z*_n_, was modeled using linear regression:

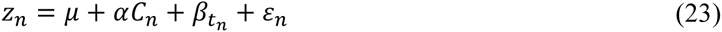

where, *C*_n_ is the *n*^th^ column of the matrix *C*_*M×N*_ containing *M* confounding factors, and *t*_n_ ∈ {0, 1, 2}, indicates the number of alternative alleles in the *n*^th^ sample. All non-significant columns, for which 95% confidence interval of the regression coefficient in α overlapped zero, were discard from *C*. In the second step, the regression was repeated using the reduced covariate matrix and corrected expression were derived as

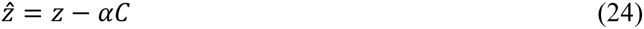

Corrected expression vector, 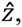 was used for effect size calculations. For direct estimation of aFC from Eq. 17 (the Nonlinear method, M2, in Fig. 3, 4), we used Matlab generic nonlinear least square solver (*lsqnonlin*). The effect size estimates used in Fig. 5, as well as those published on GTEx portal (http://gtexportal.org) were calculated using the nonlinear approximation method (M3), and the 95% confidence intervals for the aFC estimates were calculated using the bias-corrected and accelerated bootstrap (Efron 2012).

#### 5. Independent eQTL calling

Multiple independent signals for a given expression phenotype were identified by forward stepwise regression followed by a backwards selection step. The gene-level significance threshold was set to be the maximum beta-adjusted P-value (correcting for multiple-testing across the variants) over all eGenes in a given tissue. At each iteration, we performed a scan for *cis*-eQTLs using FastQTL (Ongen et al. 2016), correcting for all previously discovered variants and all standard GTEx covariates. If the beta adjusted P-value for the lead variant was not significant at the gene-level threshold, the forward stage was complete and the procedure moved on to the backward stage. If this P-value was significant, the lead variant was added to the list of discovered *cis*-eQTLs as an independent signal and the forward step moves on to the next iteration. The backwards stage consisted of testing each variant separately, controlling for all other discovered variants. To do this, for an eGene with n eVariants we ran *n cis* scans (in effect *n* − 1 *cis* scans, as one replicates the final stage of the forward analysis). For each cis scan we control for all covariates and all but one of the discovered eVariants (the one dropped is the genetic signal that is being tested, conditioned on the full model). If no variant was significant at the gene-level threshold the variant in question was dropped, otherwise the lead variant from this scan, which controls for all other signals found in the forward stage, was chosen as the variant that represents the signal best in the full model.

#### 6. Joint analysis of two eQTLs

##### 6.1 Regulatory independent model

Let us assume two biallelic eVariants, *v*_1_ and *v*_2_ regulating expression of the same eGene in *cis*. This is a special case of Eq. 5-7 where *N*=2, and *m*_1_=*m*_2_=2. Under independence assumption, regulatory effect of each eVariant allele on the expression of the carrying haplotype does not depend on the present allele for the other eVariant, and therefor, the expression of a haplotype carrying alleles *i*_1_ and *i*_2_ for the two eVariants is

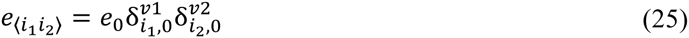

where indices *i*_1_, *i*_2_ ∈ {0,1} indicate reference (zero) and the alternative allele (one), and 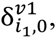 and 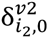 are the aFCs associated with the present alleles relative to the reference allele, for *v*_1_ and *v*_2_, respectively, and *e*_0_ is the expression of a haplotype carrying reference allele for both eVariants. Under this model the log ratio between the expressions of the two haplotypes is

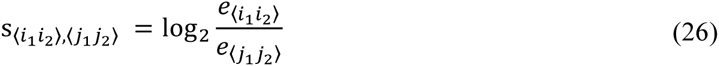

where indices *i*_1_, *i*_2_ ∈ {0,1} and *j*_1_, *j*_2_ ∈ {0,1} indicate the present alleles on the first, and on the second haplotype, respectively. From definition of aFC

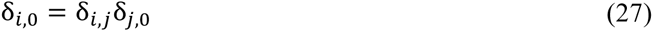

thus, after substituting haplotypic expressions from Eq. 25 in Eq. 26, the log ratio between the expressions of the two haplotypes is

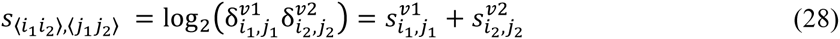

This equation presents the expected log aFC for a given genotype. Therefore, under regulatory independence model, joint effect of the two alternative alleles is sum of their individual effects:

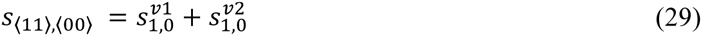

Under the *cis*-regulatory model, total expression of the eGene for each genotype is the some of the individual haplotype expressions:

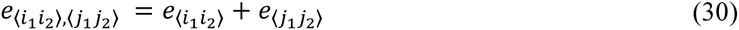

Substituting Haplotypic expressions from Eq. 25, we can use measured expression profiles of genotyped individuals to estimate aFC associated with the two eVariants. Observed expression value for the eGene in the *n*^th^ sample after log transformation is

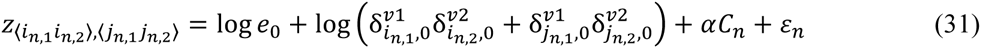

where indices *i*_*n*,1_, *i*_*n*,2_, *j*_*n*,1_, *j*_*n*,2_ ∈ {0,1} indicate the present alleles, and *C*_n_ is the provided column vector of the confounding factors for the sample. The nonlinear regression problem can be solved to estimate reference expression *e*_0_, individual aFC effects 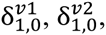 and the cofactor weight vector α (By definition 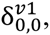 and 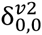 are equal to 1).

In order to estimate aFCs for eGenes with two eQTLs in GTEx data, we used PEER (Stegle et al. 2012) and top three principal components of the genotype matrix as the confounding factors in matrix *C*. Generic nonlinear least square optimizer in Matlab (*lsqnonlin*) was used to derive parameter estimates for the Eq. 26 regression problem. Confidence intervals of the parameters were derived using the *t*-statistic estimated via Jacobean matrix calculated at the optimal function values (Matlab function: *nlparci*). Predicted aFCs for regulatory independence model presented in Fig. 6B-E, and Fig. S2C (blue bars) were derived using Eq. 28.

##### 6.2 Relaxed model

In this model we relax the regulatory independence assumption, allowing the regulatory effect associated with co-occurrence of the two alternative alleles to be potentially different from sum of their individual effects. In contrast to Eq. 25, haplotype expression is

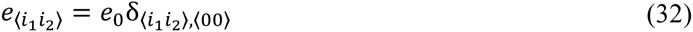

where, δ_〈*i*_1_*i*_2_〉, 〈00〉_ is the aFC associated to co-presence of the alleles *i*_1_ and *i*_2_ of the eVariants *v*_1_ and *v*_2_ as compared to a haplotype carrying reference allele for both eVariants. This model is equivalent to a special case of models in Eq. 5-7 where *N*=1, and *m*_1_=4. From aFC definition

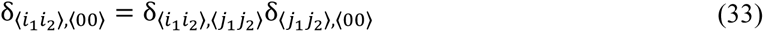

and the log ratio between the expressions of the two haplotypes is

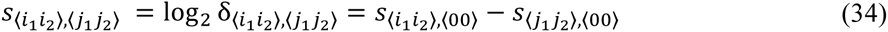

Total expression is the sum of the individual haplotypic expressions (Eq. 30), thus, the observed expression value for the eGene in the *n*^th^ sample under the relaxed regulatory model after log transformation is

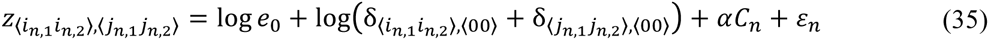

where indices *i*_n,1_, *i*_n,2_, *j*_n,1_, *j*_n,2_ indicate the present alleles and *C*_n_ the covariates as described in Eq. 31. The nonlinear regression problem can be solved for reference expression *e*_0_, joint aFC effects δ_〈10〉,〈00〉_, δ_〈01〉,〈00〉_, δ_〈11〉,〈00〉_, and the cofactor weight vector α (By definition δ_〈00〉,〈00〉_, is equal to 1).

To estimate aFCs in GTEx data, regression parameters and their confidence intervals were estimated as described for the regulatory independence model. Predicted aFCs for the relaxed model presented in Fig. 6E, and Fig. S2C (red bars) were derived using Eq. 34.

##### 6.3 Model comparison

In order to compare the two models of *cis*-regulation, the independence and the relaxed model, we calculated total data likelihood for each of the models under log-normality assumption:

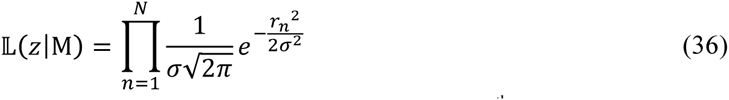

where, 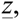 is the vector of *N* samples, and *r*_n_ is the fit residual at the *n*^th^ sample using the model considered M, and *σ* is the standard deviation of the fit residuals. Bayesian information criterion (BIC) for each of two models was calculated:

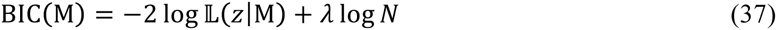

where λ, the number of parameters in each model, is the number of cofactor coefficients plus 3 and plus 4 for the regulatory independence, and the relaxed model, respectively. We used bias-corrected and accelerated bootstrap (Efron 2012) to estimate confidence intervals for ΔBIC = BIC(Relaxed model) – BIC(Independence model) in cases where ΔBIC negative. The relaxed model was selected in cases were the upper bound for the 95% confidence interval for ΔBIC fell below zero, and for the rest of the cases the independence model that has fewer parameters was deemed adequate. Calculated aFCs for all eGenes in GTEx with two associated eQTLs are provided in **Table S1**.

### Data access

The full data of the GTEx V6p release are available in dbGaP (study accession phs000424.v6.p1), and eQTL summary statistics, including the effect size estimates for the top eVariant–eGene pair per tissue [to be released at publication], are available from the GTEx Portal (http://gtexportal.org). Software for calculating allelic fold change from standard eQTL data is available in GitHub (https://github.com/secastel/aFC).

## Acknowledgments

The Genotype-Tissue Expression (GTEx) Project was supported by the Common Fund of the Office of the Director of the National Institutes of Health. Additional funds were provided by the NCI, NHGRI, NHLBI, NIDA, NIMH, and NINDS. Donors were enrolled at Biospecimen Source Sites funded by NCI\SAIC-Frederick, Inc. (SAIC-F) subcontracts to the National Disease Research Interchange (10XS170), Roswell Park Cancer Institute (10XS171), and Science Care, Inc. (X10S172). The Laboratory, Data Analysis, and Coordinating Center (LDACC) was funded through a contract (HHSN268201000029C) to The Broad Institute, Inc. Biorepository operations were funded through an SAIC-F subcontract to Van Andel Institute (10ST1035). Additional data repository and project management were provided by SAIC-F (HHSN261200800001E). The Brain Bank was supported by a supplements to University of Miami grants DA006227 & DA033684 and to contract N01MH000028. Statistical Methods development grants were made to the University of Geneva (MH090941 & MH101814), the University of Chicago (MH090951, MH090937, MH101820, MH101825), the University of North Carolina - Chapel Hill (MH090936 & MH101819), Harvard University (MH090948), Stanford University (MH101782), Washington University St Louis (MH101810), and the University of Pennsylvania (MH101822). The data used for the analyses described in this manuscript were obtained from dbGaP accession number phs000424.v6.p1 on 05/23/2016. TL and PM are supported by NIH grant R01MH106842, TL is supported by the NIH grant UM1HG008901, and TL and SEC are supported by the NIH contract HHSN2682010000029C and R01MH101814. The multiple eQTL mapping was performed at the Vital-IT(http://www.vital-it.ch) Center for high-performance computing of the SIB Swiss Institute of Bioinformatics.

### Author contributions

P.M. and T.L. designed the study. P.M. developed the statistical models and the MATLAB toolbox, and analyzed the data. S.E.C. developed the Python package and analyzed the data. A.A.B provided the independent eQTL data. P.M. and T.L. wrote the manuscript with contributions from all the authors. All the authors read and approved the final manuscript.

### Disclosure declaration

The authors declare no competing financial interests.

## Supplemental methods

### Derivations and proofs

1. **Proof: Log-transformed eGene expression is linear in the number of alternative alleles as the *cis*-regulatory effect size approaches zero**. Let *α*_1_ and *α*_2_ be the slope of the line connecting eGene expressions from reference homozygous to heterozygous, and from heterozygous to homozygous alternative genotype, respectively, in the piecewise linear model of log-transformed eQTL data (Fig. 1C):

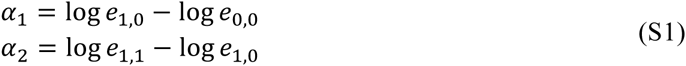 In a linear model *α*_1_ is equal to *α*_2_. Substituting the allelic expressions from the main text Eq. 4, the ratio between the two slopes for weak eQTLs is

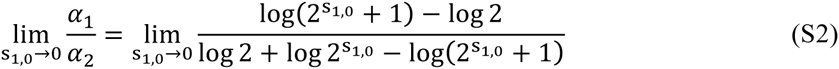

where, *s*_1,0_ = log_2_ δ_1,0_ is the eQTL effect size. Since, the limit value for both nominator and the denominator is 0, we apply L’Hôpital’s rule

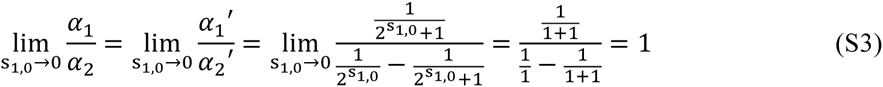 Thus, the two slopes, *α*_1_ and *α*_2_ are equal in weak eQTLs as s_1,0_ → 0.
2. **Derivations: Approximate nonlinear model for aFC estimation** Let us assume *t*_n_ is the number of alternative allele in *n*^th^ sample, and *m*_0_, *m*_1_, and *m*_2_ are the geometric means of expression in the samples homozygous for reference allele (*t*_*n*_ = 0), heterozygous (*t*_*n*_ = 1), and homozygous for the alternative allele (*t*_*n*_ = 2) respectively. First, we use the expression ratio between each of the two genotype classes to estimate aFC. From Eq. 17, the expected log-transformed expression at each eQTL genotype class is

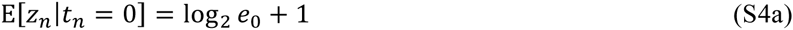

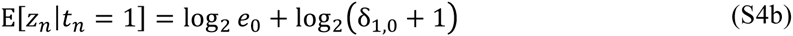

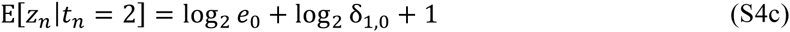 Using Eqs. S4a, and S4c, the log_2_ aFC is

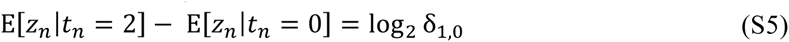 Substituting observed geometric means *m*_*t*_ = 2^E[*z*_*n*_|*t*_*n*_ =*t*]^, and exponentiating both sides of the equation, the aFC is

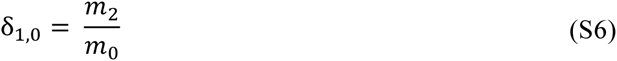 Next, we use Eqs. S4b, and S4c:

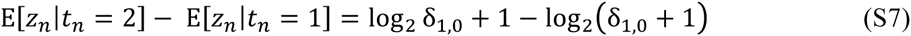 Exponentiating the both sides we have

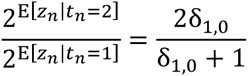

after substituting geometric means and rearranging the terms, the aFC is given:

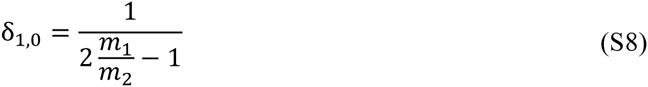 Using Eqs. S4a, and S4b

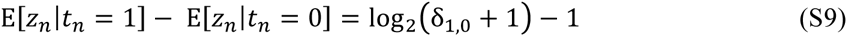

aFC can be similarly derived:

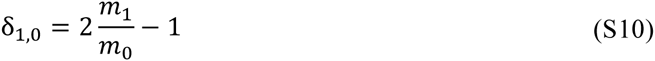 As a fourth estimate, we use loglinear regression to derive another aFC estimate. This is an accurate model for weak eQTLs where the piece-wise linear eQTL model approaches linearity (see Eqs. S1–3). The regression line passes E[*z*_*n*_ |*t*_*n*_ = 0] at *t*_*n*_= 0, and E[*z*_*n*_ |*t*_*n*_ = 2] at *t*_*n*_= 2, therefore the slope, *c*_1_, of the line is

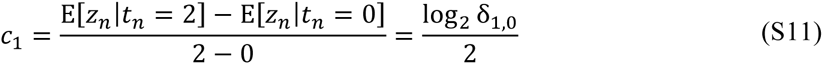 Thus aFC is given as

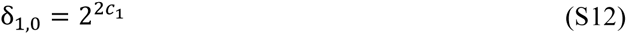 It is worth noting that under the *cis*-regulatory model of Eqs. 4a-c, the expression in the heterozygous class is at least half of that of the higher expressed homozygous class, taking place when the weak allele is effectively zero expressed, thus:

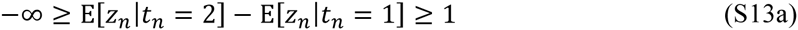

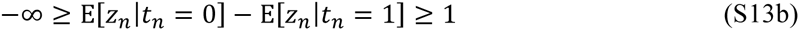 In practice, the observed expression of the genotype classes, *m*_0_, *m*_1_, and *m*_2,_ can occasionally fall outside these boundaries due to noise or other confounding biological factors beyond the considered *cis*-regulatory model. Therefore, the ratios 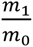 and 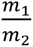 in Eqs. S8 and S10 should be bound to be ≥0.5 to avoid negative aFC estimates.
3. **Mathematical properties of log aFC** Recalling log aFC definition:

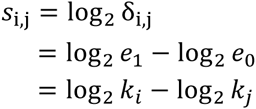 We show that the following statements are true:
  a. **Zero log aFC indicates the absence of regulatory difference: *s*_*i,i*_ = 0**

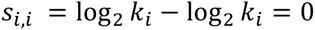
  b. **Choice of reference allele only affects the sign of log aFC: *s*_*i,j*_ = −*s*_*j,i*_**

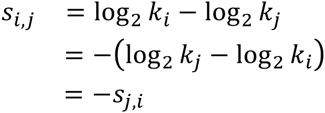
  c. **Log aFC is additive: *s*_*i,k*_ = *s*_*i,j*_ + *s*_*j,k*_**

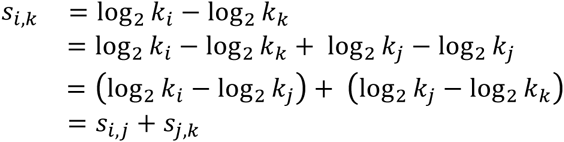
  d. **aFC associated with joint effect of independent regulatory variants, *v*1…*v*N is sum of their individual log aFCs**:

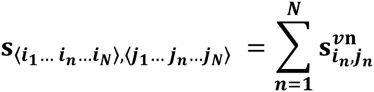

**where 〈*i*_1_… *i*_n_… *i*_N_〉 and 〈*j*_1_… *j*_n_… *j*_N_〉 are the set of present alleles on each of the haplotypes**. Assuming that variants affect gene expression independently, haplotype expression in Eq. 1 in the main text can be written for *N* eVariants as

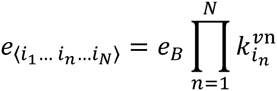

where 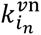 denotes the regulatory effect on the eGene expression specific to allele *i*_*n*_ of the *n*^th^ eVariant. Therefore, the joint aFC is

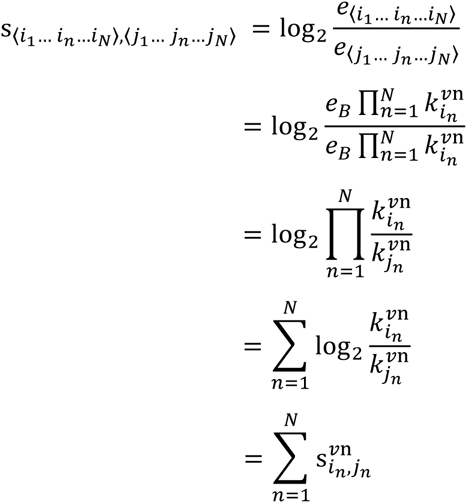
  e. **Absolute value of log aFC, *d*_*i,j*_ = |*s_i,j_|, is a pseudo-metric***:
    i. ***d*_*i,j*_ ≥ 0**
    ii. ***d*_*i,i*_ = 0**
    iii. ***d*_*i,j*_ = *d*_*j,i*_**
    iv. ***d*_*i,k*_ ≤ *d*_*i,j*_ + *d*_*j,k*_**
  The first condition is met by definition and the second and third conditions are trivial considering the aFC properties **S.I** and **S.II** shown above. In order to demonstrate the truth of the fourth condition we consider two cases:

1. When *s*_*i,j*_ and *s*_*j,k*_ are both positive or both negative; in such cases due to additivity of log aFC (Statement **S.III**), *s*_*i,k*_ will also have the same sign, and therefore, *d*_*i,k*_ = *d*_*i,j*_+ *d*_*j,k*_ is trivial.
2. When *s*_*i,j*_ and *s*_*j,k*_ have different signs; Let us assume *s*_*i,j*_ ≥ 0 and *s*_*j,k*_ ≤ 0, from **S.III**:

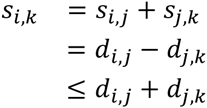 Additionally,

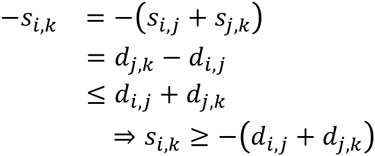 Combining the last two statements *d*_*i,k*_ = |*s*_*i,k*_| ≤ −*d*_*i,j*_ + *d*_*j,k*_. The opposite case where *s*_*i,j*_ ≤ 0 and *s*_*j,k*_ ≥ 0, is the same.

**Figure S1:**
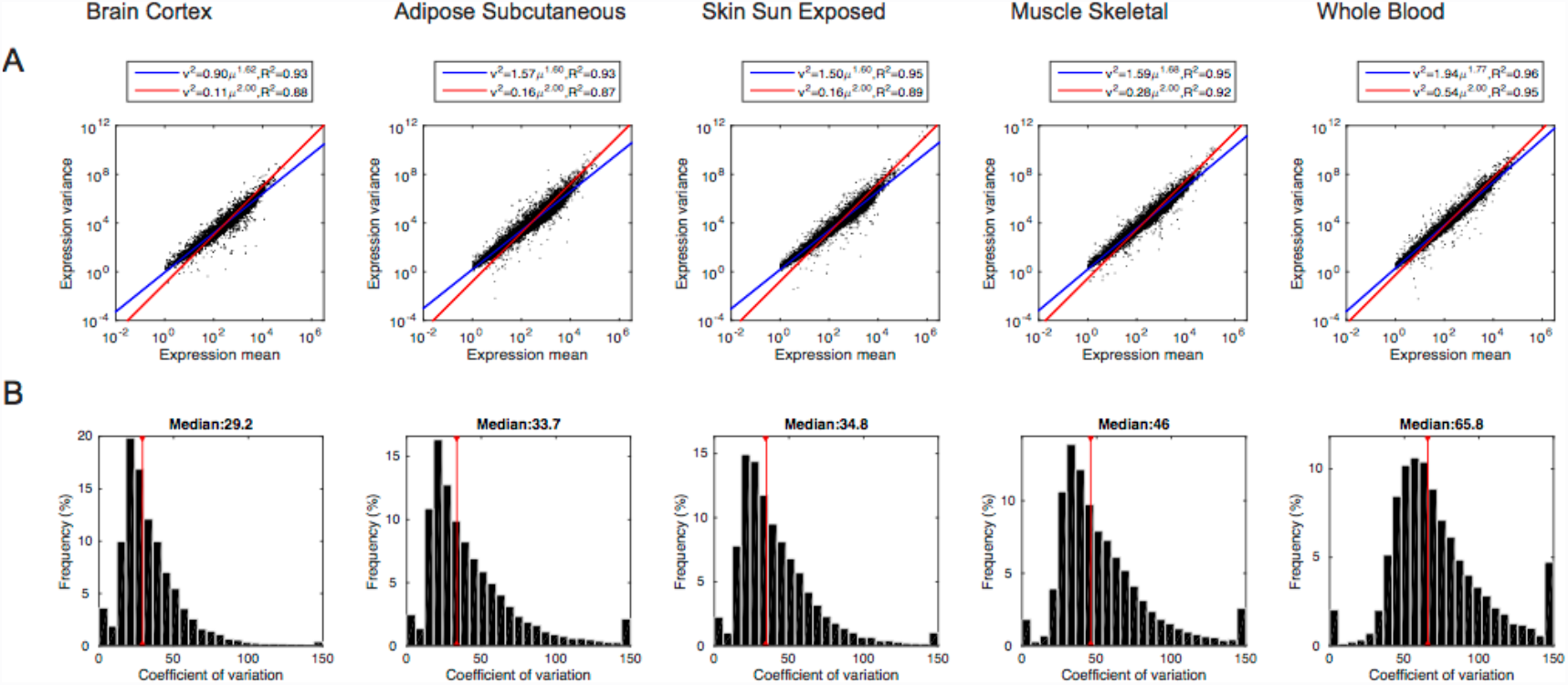
Gene expression noise distribution in GTEx data. A) Mean and variance of eGene expression within genotype classes of the top eQTL for five example tissues in GTEx data. Each dot corresponds to data from one eGene within an eQTL genotype class. Red line indicates the best expected mean-variance dependence from lognormal distributed data, and blue lines shows the optimal linear regression line. This pattern shows that variance structure eQTL data is highly similar to lognormal distribution. B) Coefficient of variation, the ratio between the standard deviation and mean, for eGene expression within eQTL genotype classes for the same tissues.

**Figure S2:**
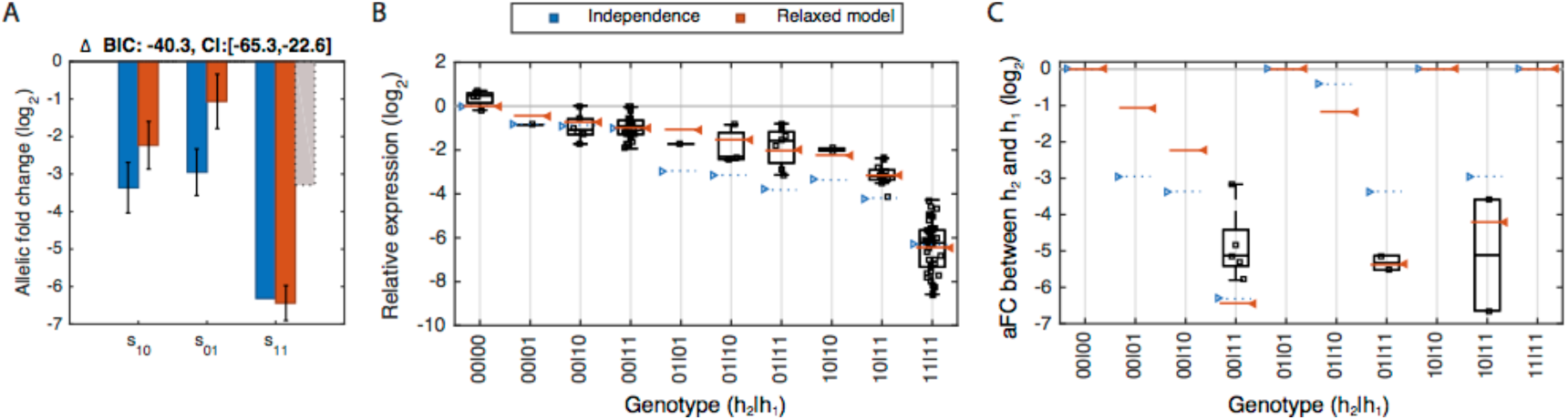
An example of two eQTLs regulating the expression of the same gene that is not well described by their individual regulatory effects acting independently (eGene: *HLADQB1-AS1*; eVariant1: chr6:32627082 A/G and eVariant2: chr6:32609813 T/C; Tissue: LCL) A). Estimated log aFC associated with the alternative alleles for the first (*s*_10_), and the second eSNP (*s*_01_) individually, along with the estimated log aFC associated with co-occurrence of the alternative alleles (*s*_11_). The independent regulation model is shown in blue, where *s*_10_, *s*_01_ and *s*_11_ are estimated from the data with the constraint of *s*_11_ = *s*_10_ + *s*_01_. The red bars show estimates from the alternative, relaxed model which allows for non-independence or epistatic-like interaction between the two eVariants, and *s*_10_, *s*_01_ and *s*_11_ are estimated without assuming *s*_11_ = *s*_10_ + *s*_01_. The support for non-independent effects comes from the difference between this estimated *s*_11_ to the sum of *s*_10_ and *s*_01_ from the relaxed model (gray dashed bar), which represents the expected joint effect of the two alternative alleles had they acted independently. B) Relative expression of the eGene and the model fits for the different genotype classes. Each dot is the expression observed in one individual, and expression levels are shown relative to the all-reference genotype. The blue and red bars show best fits achieved with and without the regulatory independence assumption, respectively. The model assuming regulatory independence between the two eVariants fails to adequately describe the observed data as measured by significance of BIC difference. C) Expression of the second haplotype relative to the first haplotype shown for different genotype groups. The dots indicate the observed values in ASE data and the blue and red bars show predicted values from the model fitted on eQTL data (as shown in panel B) using regulatory independence and the relaxed model, respectively. Genotypes in panel B and C are labeled following the notation in Fig. 6, and classes identical with regard to the *cis*-regulatory model are collapsed together in each panel.

